# The relationship between eDNA particle concentration and organism abundance in nature is strengthened by allometric scaling

**DOI:** 10.1101/2020.01.18.908251

**Authors:** M.C Yates, D. Glaser, J. Post, M.E. Cristescu, D.J. Fraser, A.M. Derry

## Abstract

Organism abundance is a critical parameter in ecology, but its estimation is often challenging. Approaches utilizing eDNA to indirectly estimate abundance have recently generated substantial interest. However, preliminary correlations observed between eDNA concentration and abundance in nature are typically moderate in strength with significant unexplained variation. Here we apply a novel approach to integrate allometric scaling coefficients into models of eDNA concentration and organism abundance. We hypothesize that eDNA particle production scales non-linearly with mass, with scaling coefficients < 1. Wild populations often exhibit substantial variation in individual body size distributions; we therefore predict that the distribution of mass across individuals within a population will influence population-level eDNA production rates. To test our hypothesis, we collected standardized body size distribution and mark-recapture abundance data using whole-lake experiments involving nine populations of brook trout. We correlated eDNA concentration with three metrics of abundance: density (individuals/ha), biomass (kg/ha), and allometrically scaled mass (ASM) (∑(individual mass^0.73^)/ha). Density and biomass were both significantly positively correlated with eDNA concentration (adj. R^2^ = 0.59 and 0.63, respectively), but ASM exhibited improved model fit (adj. R^2^ = 0.78). We also demonstrate how estimates of ASM derived from eDNA samples in ‘unknown’ systems can be converted to biomass or density estimates with additional size structure data. Future experiments should empirically validate allometric scaling coefficients for eDNA production, particularly where substantial intraspecific size distribution variation exists. Incorporating allometric scaling may improve predictive models to the extent that eDNA concentration may become a reliable indicator of abundance in nature.

## Introduction

Developing methods to estimate animal abundance in nature has attracted the attention of researchers and managers alike for over a century (Schwarz & Seber, 1999). Abundance is a fundamental population parameter in ecology, conservation, and natural resource management (Luikart, Ryman, Tallmon, Schwartz, & Allendorf, 2010), with direct impacts on ecological interactions (Krebs, 2009), ecosystem functioning (Schaus et al., 2010), population persistence and adaptability (Jamieson & Allendorf, 2012), as well as ecosystem services/resources (Immell & Anthony, 2008; Schwarz & Seber, 1999). Methodologies to estimate animal abundance represent a well-developed field of empirical research in ecology that has progressed remarkably (Schwarz & Seber, 1999; Seber, 1986). Yet despite this success, the estimation of abundance in nature is often challenging; obtaining robust estimates in natural populations using traditional methods can be time-consuming, costly, labor intensive, or even impossible to obtain for some populations (Luikart et al., 2010; Ovenden et al., 2016; Yates, Bernos, & Fraser, 2017).

The recent development of novel molecular tools has renewed interest in utilizing genetic information to indirectly estimate abundance in difficult-to-sample natural populations (Goldberg, Strickler, & Pilliod, 2015; Luikart et al., 2010). Molecular techniques that quantify the concentration of environmental DNA (eDNA) particles represent a promising tool, with recent studies demonstrating support for a correlation between eDNA concentration and abundance (Pilliod, Goldberg, Arkle, & Waits, 2013; Takahara, Minamoto, Yamanaka, Doi, & Kawabata, 2012; Thomsen et al., 2012). In addition to monitoring of species of conservation concern, eDNA represents a potential indirect-but-accurate means to quantify abundance that has broad implications for species harvesting, invasive species control, and monitoring of key indicator species used to assess ecosystem health (Barnes & Turner, 2016).

Laboratory studies have demonstrated a strong correlation between eDNA concentration and abundance (Eichmiller, Miller, & Sorensen, 2016; Klymus, Richter, Chapman, & Paukert, 2015), exhibiting a mean correlation coefficient of 0.9 (R^2^ = 0.81) (Yates, Fraser, & Derry, 2019). Studies in nature, however, have generally found weaker correlations than laboratory studies, with a mean correlation coefficient of 0.71-0.75 (R^2^ = 0.51-0.57) (Yates et al., 2019). Although correlations remain moderately strong in nature, much of the variation in eDNA particle concentration across environments often remains unexplained. As a result, the extent to which eDNA could be used to reliably infer abundance in nature remains limited without significant improvements in modelling or technology.

In nature, organismal abundance is typically quantified by evaluating individual density (i.e. individuals/unit area) or biomass density (i.e. kg/unit area). While both metrics of abundance appear to correlate equally well with species-specific eDNA particle concentration in the wild, processes involved in the production of eDNA particles in natural environments are unlikely to scale linearly with either biomass or density. Although eDNA production tends to increase with individual mass (Maruyama, Nakamura, Yamanaka, Kondoh, & Minamoto, 2014), individuals with a large biomass often produce fewer eDNA particles than equivalent biomass of smaller conspecifics (Maruyama et al., 2014; Mizumoto, Urabe, Kanbe, Fukushima, & Araki, 2018; Takeuchi, Iijima, Kakuzen, Watanab, & Yamada, 2019). As such, eDNA particle concentration would be expected to vary, for example, between environments that contain equal densities of individuals but with varying biomass. Similarly, environments with equal biomass but varying densities would also be likely to vary in observed eDNA particle concentration. Wild populations often exhibit substantial inter-population variation in the distribution of individual biomass (Donald, Anderson, Mayhood, Anderson, & Correlations, 1980; Guernon, Yates, Fraser, & Derry, 2018; Millien et al., 2006; Sebens, 1987), which may in turn scale to affect overall population-level rates of eDNA production (Maruyama et al., 2014) and partially account for the substantial unexplained variation observed between eDNA concentration and traditional metrics of abundance (e.g. density and biomass) in nature (Yates et al., 2019).

Here, we extend models of physiological allometric scaling to organismal eDNA particle production to provide a framework through which differences in density, total biomass, and the distribution of individual biomass can be integrated into models of eDNA production in natural populations. Allometry refers to changes in organisms (e.g. physiological rates, morphology, etc.) that occur in relation to proportional changes in body size (Gittleman, 2011). Excretory processes (urine, fecal matter, etc.) and shedding (from scales, skin, mucous, etc.) are thought to be the two major physiological processes that contribute to the production of eDNA particles (Jo, Murakami, Yamamoto, Masuda, & Minamoto, 2019; Stewart, 2019). The metabolic theory of ecology (MTE) provides a robust, empirically validated framework through which allometry in metabolic processes (including excretion) can be modelled (Brown, Gillooly, Allen, Savage, & West, 2004). The MTE posits that metabolic processes scale non-linearly with body size according to the power function:

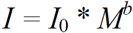

where *I* = metabolic rate, *I*_0_ = a normalization constant, *M* = organism body mass, and *b* = an allometric scaling coefficient (Allegier, Wenger, Rosemond, Schindler, & Layman, 2015; Brown et al., 2004; Vanni & McIntyre, 2016). The value of *b* varies depending on the physiological process; metabolic rates typically scale to the power of 0.75 (Brown et al., 2004; Isaac & Carbone, 2010), whereas values for consumptive or excretory rates are often lower (Post, Parkinson, & Johnston, 1999; Vanni & McIntyre, 2016). Nevertheless, metabolic theory predicts that larger organisms tend to exhibit disproportionately lower rates (relative to their mass) for metabolically linked processes such as excretion (Allen & Gillooly, 2009; Vanni & McIntyre, 2016). While shedding from mucous, scales, or skin may also be linked to metabolic rates, shedding rates are also likely a function of the surface area of an organism. In many aquatic organisms (particularly fish) the allometric relationship between body mass and surface area follows a similar mathematical form as metabolic processes; salmonids, for example, exhibit mass-scaling coefficients for surface area between 0.59 and 0.65 (Shea, Fryer, Pert, & Bricknell, 2006).

Metabolic rates, excretory rates, and surface area (via shedding) are likely to collectively impact eDNA production, yet all follow a similar allometric form. As a result, we hypothesize that eDNA production can also be modelled as an approximate power function of individual mass and an exponential scaling coefficient with a value less than 1. That is, the rate at which eDNA production increases with body mass will decline (Figure 1a) such that, on a per-gram basis (e.g. mass-specific rate), large individuals will tend to excrete fewer eDNA particles relative to smaller conspecifics (Figure 1b). This hypothesis has important consequences for ecosystem-level processes; the utility of integrating allometric scaling in ecosystem-level models of ecological stoichiometry (Allen & Gillooly, 2009), animal excretion (Vanni & McIntyre, 2016), consumption (Post et al., 1999), and nutrient cycling (Schaus et al., 2010; Schindler & Eby, 1997), for example, has long been acknowledged with broad empirical support. We therefore further hypothesize that, when scaled to the level of an entire population, allometric scaling in eDNA production will also have a substantial effect on overall population-level production of eDNA. We consequently predict that the incorporation of mass scaling coefficients to account for inter-population variation in density, biomass, and the distribution of biomass across individuals will improve modelling efforts linking eDNA particle concentration and abundance across natural ecosystems.

**Figure 1:**
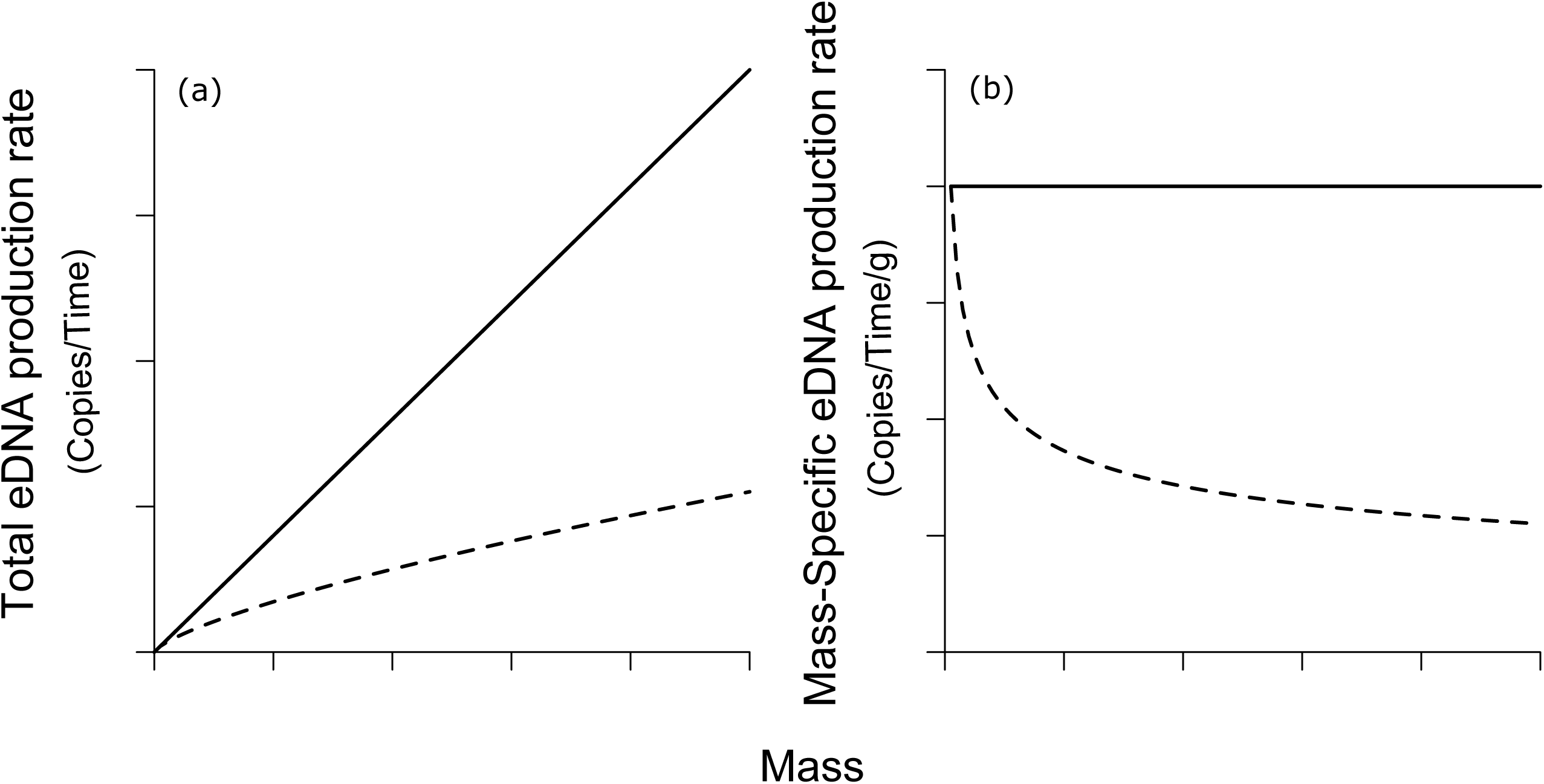
Conceptual example of allometric scaling in eDNA production rate with individual mass. Figure a) demonstrates absolute eDNA production rate as mass increases, figure b) demonstrates mass-specific eDNA production rate (i.e. eDNA production rate per (g) of mass). Solid lines reflect an allometric process that scales linearly with body mass (*b* = 1), dashed lines correspond to allometric scaling with a value of *b* < 1.

To test our hypothesis, we collected standardized individual biomass data and used common mark-recapture experiments to enumerate abundance in nine populations of brook trout (*Salvelinus fontinalis*) in the Rocky Mountains of Canada while simultaneously collecting eDNA samples in each lake. Study populations exhibited substantial variation in individual density (63 - 1177 individuals/ha), biomass density (12.6 - 52.4 kg/ha), and mean body size (43.0 - 405.9 g/individual). We applied these data to specifically test two key predictions: i) brook trout eDNA particle concentration will correlate with traditional metrics of abundance (density and biomass) across the nine study lakes; and ii) incorporating allometric scaling coefficients to estimates of brook trout abundance (e.g. ∑(individual biomass^0.73^)/ha, or “allometrically scaled mass” (ASM)) will substantially improve models of abundance and eDNA particle concentration.

ASM estimates derived from known eDNA concentrations in novel systems lacking abundance data cannot be directly converted to traditional metrics of abundance (e.g. density and biomass) because multiple density/biomass configurations (e.g. many small fish or a small number of large fish) can produce equivalent ASM values. However, using a real-world example, we also demonstrate how ASM estimates derived from known eDNA concentrations for systems that lack abundance data on a target species can be converted into traditional estimates of abundance with additional size structure data.

## Materials and Methods

### Study species and system

Nine brook trout populations introduced in the early 20^th^ century to lakes located in Kootenay, Banff, and Yoho national parks (Figure S1) were monitored to determine population size and individual biomass distributions. Brook trout represent ideal populations to study allometry in eDNA production and its impact on the relationship between eDNA particle concentration and abundance. Several studies have already demonstrated significant correlations between abundance and eDNA concentration for brook trout in lotic systems (Baldigo, Sporn, George, & Ball, 2017; Wilcox et al., 2016). Brook trout populations also often exhibit substantial variation in size structure (Donald et al., 1980; Guernon et al., 2018), providing the opportunity to study populations that represent a gradient of small-to-large bodied individuals. Additionally, our study populations experience little recreational fishing pressure due to no-take policies implemented within the National Parks.

### Mark-recapture surveys and size structure estimates

Mark-recapture studies were conducted in 2018 between May 27^th^ and June 30^th^, except for Cobb lake where isolated marking events occurred until September 12^th^ (Figure S2). Fish were captured using a combination of fyke nets, angling, and backpack electrofishing (Table 1). Large (1 m hoop diameter, 2 cm mesh) and small (0.7 m hoop diameter and 0.8 cm mesh) fyke nets were distributed around the perimeter of lakes with the lead attached to shore and the end of the trap facing the center of the lake. Nets were checked daily to reduce stress to fish and possible cannibalism. Angling was used to supplement fish capture efforts at sites where fyke catchability was low (predominantly Cobb). Marks were also assigned to fish captured by electrofishing the shore and inlets/outlets of lakes with a backpack electrofisher (Smith-Root, Vancouver, Washington, USA)

**Table 1:**
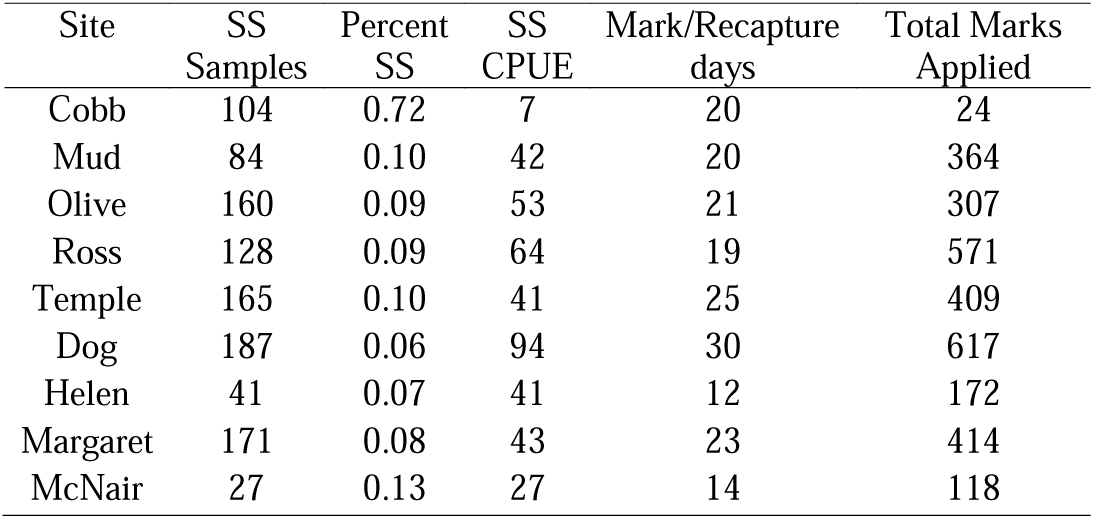
Size structure gill net effort and fyking (mark-recapture) effort. SS refers to size structure assessment, percent SS refers to the proportion of population harvested during size structure assessment.

| Site | SS<br>Samples | Percent<br>SS | SS<br>CPUE | Mark/Recapture<br>days | Total Marks<br>Applied |
| --- | --- | --- | --- | --- | --- |
| Cobb | 104 | 0.72 | 7 | 20 | 24 |
| Mud | 84 | 0.10 | 42 | 20 | 364 |
| Olive | 160 | 0.09 | 53 | 21 | 307 |
| Ross | 128 | 0.09 | 64 | 19 | 571 |
| Temple | 165 | 0.10 | 41 | 25 | 409 |
| Dog | 187 | 0.06 | 94 | 30 | 617 |
| Helen | 41 | 0.07 | 41 | 12 | 172 |
| Margaret | 171 | 0.08 | 43 | 23 | 414 |
| McNair | 27 | 0.13 | 27 | 14 | 118 |

Captured fish were anesthetized using clove oil and measured for fork length (± 1mm) and mass (± 0.1g). Any unmarked fish were gastrically tagged with a BioMark HPT8 pre-loaded Passive Integrated Transponder (PIT) tag (Boise, Idaho, USA). Only fish greater than or equal to 80 mm were tagged to reduce tagging mortality. The tag number of any recaptured fish was recorded. All fish were processed in the shade with aerators to avoid unnecessary stress. Recovered fish were released in the center of the lake to standardize release location and promote mixing (e.g. if released near shore, fish may have been recaptured in an adjacent net, biasing mark recapture data). Marking ceased once recapture ratios approached twenty five percent for several consecutive days in order to standardize marking efforts across all populations and to ensure that enough fish were tagged to facilitate census size (N_c_) estimates have confidence intervals within 10% to 25% of true values, following general methodologies reviewed in (Krebs, 2009).

Size structure estimates aimed to obtain a representative snapshot of the size structure of each population and were conducted between July 27^th^ and September 1^st^, with the exception of Cobb where size structure assessments continued to October 12^th^ (Figure S2). Fyke-nets were deployed in littoral zone areas extending to the centre of the lake and, as a result, size-structure assessments may be more biased towards small-medium bodied individuals (who prefer littoral habitats) (Tiberti et al., 2017). To obtain a relatively unbiased estimate of population size structure, fish were captured in large and small sinking mixed mesh gillnets with clear monofilament distributed throughout the lake. Large mixed-mesh gillnets were 15.6 m long, 1.8 m deep and had an equal area of 64-51-89-38-76 mm mesh panels. Small mixed-mesh gillnets were 12.5 meters long, 1.8 meters deep, and consisted of an equal area of 32-19-38-13-25 mm mesh panels. Index nets are widely used in North America for size structure assessments (Bonar, Hubert, & Willis, 2009; Hubert, Pope, & Dettmers, 2012; Johnson, 1983; Post et al., 1999; Ward, Askey, Post, Varkey, & Mcallister, 2012) as these attempt to capture a representative size/age structure of the population (Morgan, 2002). Nets were checked daily and moved to different locations across the lake if reset in order to capture a representative sample of fish in each lake. Sampling ceased when approximately five to ten percent of the population was captured, apart from Cobb lake where size structure assessment captured approximately 71% of individuals (Table 1). Captured fish were euthanized with clove oil, PIT tags were recorded, and length/mass data were collected as described for the marking period.

### Population size estimation

Schnabel population size estimates, which utilize sequential marking/recapture events, were used to determine the number of fish in a lake (Schnabel, 1938). All size structure assessment removals were pooled together into one final sampling event for the population estimates which controlled for the removal of marks at large. Note that population estimates only account for fish greater than the minimum tagging size (80 mm fork length). All population estimates were conducted in R (R Development Core Team, 2017) with the *mrClosed* function from the Fisheries Stock Assessment package FSA (Ogle, 2016). Confidence intervals for Schnabel population estimates followed recommendations from (Seber, 2002) as implemented in the FSA package.

### Density calculation

To link eDNA particle concentration with fish abundance, three metrics of density were calculated: (i) individual density (individuals/ha); (ii) biomass density (biomass/ha); (iii) and allometrically scaled mass (ASM/ha). Individual density was estimated by dividing the population size estimate by lake size (ha). Biomass density was calculated according to the following formula:

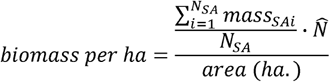

Where 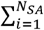 *Mass*_*SAi*_ is the sum of the masses captured in the index net during size structure assessment, N_SA_ is the number of fish captured in the index nets, 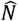 is the estimated population size. This methodology assumes that the size structure assessment was representative of the population.

ASM was calculated by replacing the mass measure with *mass*^*0.73*^ according to the formula:

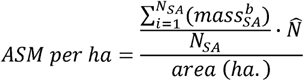

This density metric was included to account for the relative decline in mass-specific eDNA production or excretion rates typically observed as individual organismal mass increases (Maruyama et al., 2014; Takeuchi et al., 2019; Vanni & McIntyre, 2016). Scaling coefficients (the value of *b*) can vary substantially depending on the physiological process, taxonomy or environment (Allegier et al., 2015; Glazier, 2005). In the absence of data on allometric scaling in eDNA production, data on allometric scaling in metabolic or excretory rates for the same study species can represent useful starting points. Data on allometry in excretory rates were unavailable for brook trout. However, in laboratory experiments (Hartman & Cox, 2008) found that mass-specific metabolism scaled as a power law of mass with an exponent of -0.265. The scaling exponent for absolute metabolism would therefore be 1 – 0.265 = 0.735, which we used as the value of *b* in our ASM model.

In difficult to sample populations, estimates of relative abundance are often obtained using catch-per-unit-effort (CPUE) metrics. As a result, most previous studies examining eDNA particle concentration and abundance utilize similar metrics (Yates et al., 2019). To evaluate the utility of CPUE as a ‘proxy’ metric of abundance in our study system, CPUE for each lake was quantified as the mean catch per-unit effort of a large and small index gillnet.

### eDNA sample collection

eDNA samples were collected between June 30 and July 13^th^, 2018. Sampling was equidistantly distributed around each lake and included four littoral and four pelagic samples. Littoral samples were collected approximately 1-3 m from shore at a depth of least 30 cm but 15 cm above the bottom to avoid the unintentional collection of sediments, which can contain concentrated eDNA but also inhibit PCR reactions (Turner, Uy, & Everhart, 2015). Surface pelagic samples were collected from each lake along a vertical line through its center (identified as the midpoint of its longest axis); samples were collected along this axis at equidistant intervals within the first meter of depth (approximately 0.5m). eDNA for most fish species tends to be uniformly distributed throughout the water column of deep lakes (Hanfling et al., 2016) and shallow ponds (Evans et al., 2017). A thermal profile of the lake (e.g. temperature reading every 0.5 m using a YSI professional series sonde (model 10102030) (Yellow Springs Inc., Ohio, USA)) at the deepest point was taken immediately after sample collection. To avoid between-lake contamination all eDNA samples were collected either from an inflatable kayak that was decontaminated 48h prior in a 2% regular strength household bleach solution for 15 minutes (including paddle and life-jacket) or from a canoe assigned to sample a single specific lake. Water samples were collected using sterile Whirl-Pak™ bags (Uline, Ontario, Canada).

Samples were immediately filtered on the lakeshore using two chlorophyll filtering manifolds (Wildco, Florida, USA) bleached in a 30% household bleach solution for ten minutes 2-12h prior to collection. All samples were stored in the shade prior to filtration in plastic washbasins bleached with a 30% solution for ten minutes, and all filtering was conducted in the shade under a tarp. Manifolds were transported in a Polar Bear™ backpack cooler (Polar Bear Coolers, Georgia, USA) whose interior was wiped with a 30% household bleach solution for ten minutes. Manifold components were stored after bleaching and transported individually in sealed plastic zippered bags to limit contamination. Pencils and markers were also wiped with a 30% bleach solution.

One L of sample water from each site was filtered through a 0.7µm-pore glass fibre filter (GE Healthcare Life Sciences, Ontario, Canada) using a vacuum hand pump (Soil Moisture, California, USA); each vacuum pump was decontaminated between lakes by wiping with a 30% household bleach solution and resting for ten minutes. All littoral samples were filtered on one manifold and all pelagic samples were filtered on the other. Prior to filtering lake water samples, 1 L of distilled water was filtered through each manifold as a negative control. Filters were handled using two metal forceps bleached in a 30% solution for ten minutes and transported in individual bags; one forceps was used for littoral samples and another forceps was used for pelagic samples. After filtering, filters were folded and placed directly in a sterile 2 ml microcentrifuge tube filled with 700µl AL buffer (Qiagen, Maryland, USA) which was then labelled and individually sealed in a plastic zippered bag and placed in a second cooler that was decontaminated by wiping with a 30% household bleach solution and resting for ten minutes. This cooler contained two frozen freezer-gel packs decontaminated in a 30% bleach solution for ten minutes. If a filter became clogged (i.e. < 1 L of water was filtered) the final volume of water filtered was recorded and the sample was stored in buffer. Filters were immediately transported to, and stored in, a -20 □ freezer (wiped with 30% household bleach and soaked for ten minutes) at Kootenay Crossing. Filters were stored on dry ice for transportation to Montreal (driven approximately two and a half days) where they were stored in a -80 □ freezer.

### eDNA extraction and analysis

Each filter was extracted using a Qiagen DNeasy Blood and Tissue ™ kit and Qiashdredder™ spin column following a modified extraction protocol (see Appendix S1 for details). Final DNA product was eluted into 130 µl of AE buffer and stored in a clean -20 □ freezer dedicated to the sole storage of eDNA samples. To avoid contamination between lakes, extractions were conducted on batches from a single lake with a single extraction blank of 700 µL AL buffer included as an extraction control. Decontamination procedures were identical for both manifolds, so only a single negative control was extracted per lake. All extractions were conducted in an extraction room dedicated to the handling of sensitive eDNA samples. This room receives weekly cleaning with a 10% household bleach solution and is free of PCR products or high-concentration DNA. All individuals entering the extraction room were required to wear nitrile gloves, hair nets, shoe covers, and dedicated, clean lab coats. All lab surfaces were soaked with a 20% household bleach solution for ten minutes before and after extractions. PCR Clean Wipes™ (Thermo Scientific, Massachusetts, USA) were also used to decontaminate all lab surfaces and pipettes prior to and after extracting or handling eDNA samples.

The concentration of brook trout eDNA was quantified using the TaqMan minor groove assay published in (Wilcox et al., 2013), which targets a region of the brook trout cytochrome *b* mitochondrial gene. All samples were run in triplicate at a 20 µl final reaction volume on a Stratagene MX 3000P thermal cycler using Environmental Master Mix 2.0 and 5 µl of template DNA. Forward and reverse primers were included at a final concentration of 900 nM, with the probe at a final concentration of 250 nM. Each replicate was spiked with an internal positive control to test for inhibition; any replicate that exhibited inhibition (C*t* > 1 in the internal positive control) was reanalyzed with diluted template DNA at 60% concentration (3 µl template + 2 µl of ultrapure water); this was sufficient to relieve inhibition in all cases. Standard curve template DNA was composed of a synthetic Gblock™ gene fragment (IDT, Iowa, USA) of the targeted sequence. A triplicate no template control and triplicate five-point standard curve (1250 copies/µl, 250 copies/µl, 50 copies/µl, 5 copies/µl, 2 copies/µl template concentration) were included on each 96-well plate. All qPCR reaction reagents were aliquoted into single-use volumes adequate for a single plate and reactions were prepared in the dedicated eDNA room, with the exception of the standard curve replicates due to the presence of high concentration synthetic DNA fragments. Reactions were cycled with an initial hold at 95 □ for ten minutes followed by 45 cycles of 30 seconds at 95 □ and 1 min at 60 □. eDNA particle concentration at each site was determined by averaging site-specific replicates. Final mean copy number values were converted (based on total volume of water filtered per sample) to total eDNA particle concentration per 1 L of sampled water (copies/L).

### Data Analysis

Mean eDNA particle concentration (copies/L) for each lake was calculated by first averaging eDNA particle concentrations of the four littoral and four pelagic samples to obtain mean littoral eDNA concentration and mean pelagic eDNA concentration. A weighted-mean eDNA concentration for each lake was calculated by weighing the littoral and pelagic eDNA concentrations based on the fraction of total lake area each zone represented. Our study lakes varied substantially in size (1.7 to 18.5 ha); total pelagic and littoral areas were calculated for each lake using polygons on Google Earth. In the absence of detailed bathymetry data, the total area of the littoral zone (where sunlight can reach the lake bottom to support submerged macrophyte and benthic primary production (Kalff, 2001)) was calculated by including all lake surface area up to 20m from the shore, with the remaining area assigned to the pelagic zone. A distance of 20 m was chosen because, based on personal observation, we estimate that the littoral zone of the lakes extended an average of approximately 10-15 m from the shore. The concentration of eDNA near points of high concentration (i.e. high fish density or areas where fish feed) decreases rapidly, with concentrations dropping rapidly after 5-10 m (Ghosal, Eichmiller, Witthuhn, & Sorensen, 2018). A littoral zone of 20 m reflects these processes (10-15 m littoral zone + 5-10 m for diffusion). Given these assumptions, the area of the pelagic zone expressed as a fraction of the total area of a lake increases with lake size. The relative contribution of the littoral and pelagic zones to the overall mean concentration of eDNA per lake should therefore be increasingly weighted towards the pelagic eDNA concentration as lake surface area increases.

Mean lake eDNA particle concentration (copies/L) was modelled separately as a function of the three metrics of brook trout density calculated above: individual density (individuals/ha); biomass density (kg/ha); and allometrically scaled mass (ASM) (∑(individual mass^0.73^)/ha). eDNA particle concentration was included as a dependent variable in a linear regression and a separate model for each abundance metric was fitted to the observed data. Wald *F*–tests were used to evaluate the significance of fixed-effect terms, with model log-likelihood values were values used to compare model fit using the AIC criterion (Akaike, 1974) as in (Lacoursière-Roussel, Côté, Leclerc, Bernatchez, & Cadotte, 2016), assuming that models with ΔAIC > 2 exhibit significantly reduced explanatory power (Burnham & Anderson, 2002). All analyses were conducted in R (v.3.3.3) (R Development Core Team, 2017). To assess the performance of CPUE as a ‘proxy’ metric of abundance, we also examined the relationship between density and CPUE, as well as eDNA particle concentration and CPUE, using linear regression. To assess the sensitivity of the final results to the relative size of the area of the littoral zone, we ran an additional set of models in which we halved the estimated littoral area of each lake.

### Estimating density and biomass from predicted allometrically scaled mass: a case study for population management

Predicting abundance in unknown systems from known eDNA particle concentrations would require an inversion of the modelling relationship described above: abundance would be modelled as a function of eDNA particle concentration. Predicted estimates of ASM obtained from eDNA samples for systems lacking abundance data cannot be directly converted to traditional metrics of abundance (e.g. individual density or biomass density) because multiple density/biomass configurations (e.g. many small fish or a small number of large fish) can produce equivalent ASM values. However, with additional individual mass distribution data from standardized size structure data any predicted ASM point-estimates can be converted to traditional metrics. Size structure data could be exponentiated to the power of *b* (the allometric scaling coefficient) and the resulting scaled mass values nonparametrically bootstrapped until the cumulative sum of the bootstrapped values surpass the predicted ASM. Individual density could then be estimated by totalling the number of bootstrap “samples” required to surpass the predicted ASM; biomass density could then be estimated by multiplying the predicted density value by the untransformed mean of the size distribution.

As a case study, this technique was applied to data collected from Hidden Lake (Banff, Alberta, Canada). The brook trout population of Hidden Lake was targeted as part of rotenone-based removal program by Parks Canada. eDNA samples from Hidden Lake were collected in July 2018 and extracted/analyzed using the same methodology as described above. The estimated “ASM/unit area” of the lake (including 95% prediction intervals) was calculated from the linear relationship obtained from our nine study lakes. Unfortunately, standardized size structure data were unavailable; rotenone removal efforts began in August 2018 and no brook trout remain in the system. However, prior to the use of rotenone mechanical gill netting efforts were employed during brook trout removal efforts between 2011 and August 2017 (Stitt, *pers. comm.*). By 2016 netting efforts had removed most large fish from the population, and fish older than age 0+ were between 90-140mm in length (Sullivan, 2017), although it should be noted that standardized size distribution data was unavailable. Of our nine study lakes, fish from Olive lake exhibited the smallest body mass, so size structure data from this lake was utilized as a “proxy” to calculate an approximate pre-rotenone individual density and biomass density of brook trout inhabiting Hidden Lake in 2018. Bootstrap simulations to quantify individual density and biomass density utilizing the Olive size distribution and predicted ASM of Hidden Lake were run for 1000 iterations. All analyses were performed in R (R Development Core Team, 2017).

### Predicting allometric scaling coefficient for eDNA production in brook trout

Allometric scaling coefficients are likely to fall between a value of 0 and 1; notably, (∑ individual mass^0.0^)/ha is equivalent to individual density (fish/ha) and (∑individual mass^1.0^)/ha is equivalent to biomass density (kg/ha). Although we employed an allometric scaling coefficient of 0.73 in our model (based on metabolic data from brook trout), the “true” allometric scaling coefficient for eDNA production in our system was unknown. We used our data to predict the optimal value for the scaling coefficient given the observed eDNA particle concentration and biomass distribution data observed across our study lakes. To achieve this, we iteratively generated ASM values from our data using scaling coefficients ranging from 0 to 1 (increasing by intervals of 0.01) and sequentially modelled eDNA particle concentration data as a function of each ASM value. AIC values for each model were then used to evaluate model fit. If eDNA production scales allometrically according to a power function, we predict that the AIC values across models with scaling coefficients between 0 and 1 will exhibit an approximately upward parabolic distribution with a minimum best-fit value that corresponds to an “optimal” allometric scaling coefficient. According to the general rule described in (Burnham & Anderson, 2002), models within 2 ΔAIC also exhibit substantial support; we predict that the ‘true’ allometric scaling coefficient for brook trout eDNA production in nature will fall between the range of scaling coefficients that produce models within 2 AIC of the ‘best-fit’ scaling coefficient, although future experiments will be necessary to validate our predictions. To assess the sensitivity of this analysis to the estimated size of the littoral zone, this analysis was repeated for models in which we halved the estimated littoral area of each lake. All analyses were performed in R (R Development Core Team, 2017).

## Results

### Population size estimates and density

Population size estimates ranged from 145 to 3266 individuals, individual density ranged from 63 to 1131 fish/ha, biomass density ranged from 12.6 to 52.5 kg/ha, and ASM ranged from 3707 to 18600 ASM/ha (Table 2, see Figure 2 for population size structure). Estimates of catch-per-unit-effort (CPUE) did not exhibit a significant correlation with individual density (*F*_1,7_ = 0.53, *p* = 0.491, Figure S3).

**Table 2:** Density metric estimates for each population. N_c_ = population size, ASM = allometrically scaled mass. 95% confidence intervals for ‘N’ are given in brackets.

| Site | N<sub>c</sub> | Ha | Mean<br>Individual<br>Mass (g) | Fish/ha | Kg/ha | ASM/ha | eDNA<br>(copies/L) |
| --- | --- | --- | --- | --- | --- | --- | --- |
| Cobb | 145 (94, 237) | 2.3 | 404.8 | 63 (41, 103) | 25.7 (16.5, 41.7) | 5663 | 592.2 |
| Dog | 3266 (2715, 4097) | 11.5 | 184.8 | 284 (236, 356) | 52.5 (43.6, 65.8) | 13962 | 5131.1 |
| Helen | 557 (420, 755) | 2.5 | 83.9 | 225 (168, 302) | 18.8 (14.1, 25.3) | 6187 | 2445.9 |
| Margaret | 2017 (1638, 2623) | 18.0 | 112.3 | 112 (91, 146) | 12.6 (10.2, 16.4) | 3707 | 1240.4 |
| McNair | 201 (158, 276) | 1.7 | 137.3 | 121 (93, 162) | 16.6 (12.8, 22.3) | 4736 | 3050.5 |
| Mud | 860 (733, 1040) | 7.2 | 141.9 | 119 (102, 144) | 17.0 (14.4, 20.5) | 4587 | 1138.7 |
| Olive | 1877 (1459, 2628) | 1.7 | 43.1 | 1131 (858, 1546) | 48.8 (37.0, 66.6) | 18601 | 7805.1 |
| Ross | 1392 (1211, 1635) | 6.6 | 82.5 | 211 (183, 248) | 17.4 (15.1, 20.4) | 5559 | 917.4 |
| Temple | 1655 (1369, 2090) | 3.3 | 51.1 | 509 (415, 633) | 26.1 (21.2, 32.4) | 9587 | 2076.5 |

**Figure 2:**
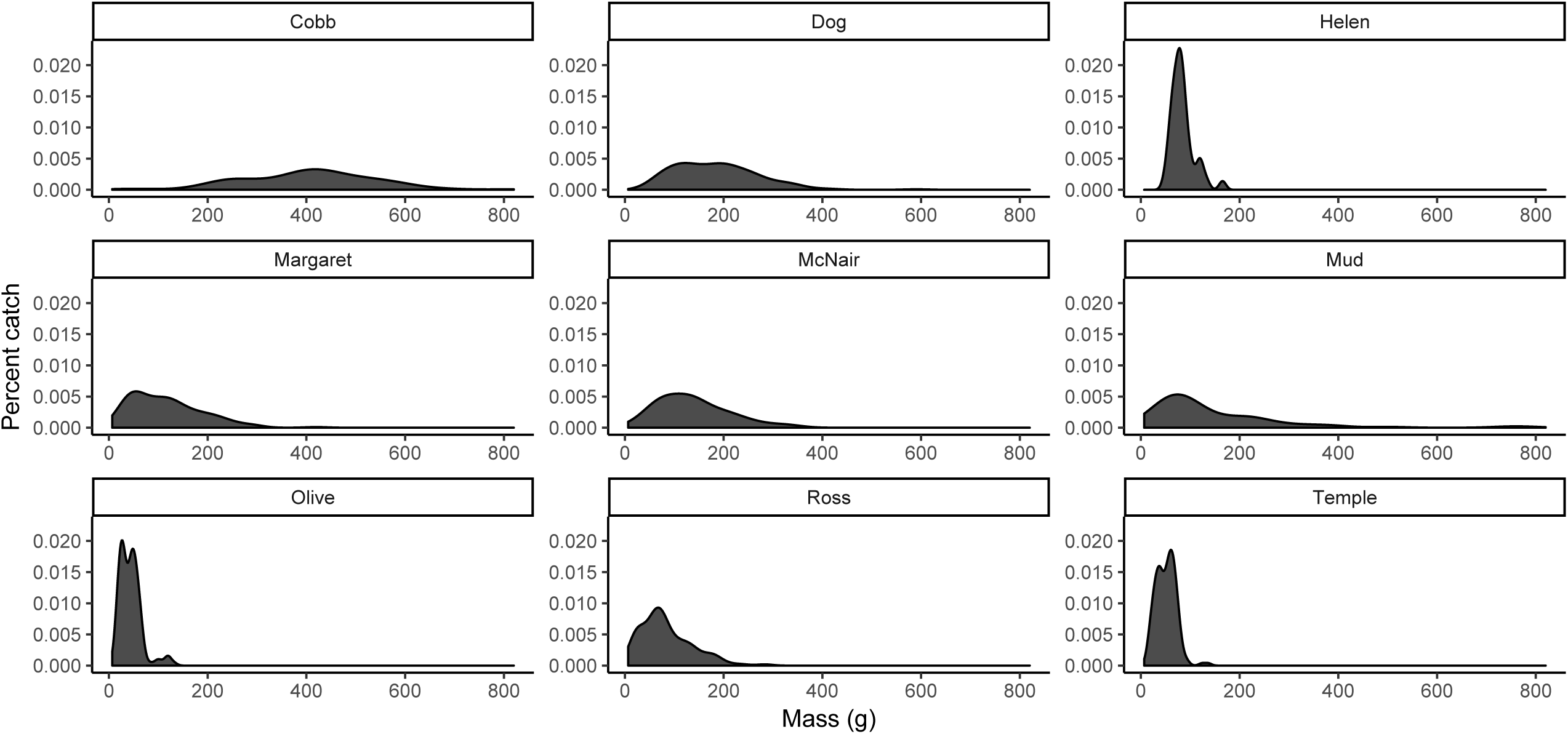
Lake size structure distributions (g) obtained from standardized gill net sets for the nine study lakes.

### eDNA concentrations and correlations with density metrics among lakes

Brook trout eDNA was successfully amplified from all samples in all lakes. No amplification was observed in any negative controls or extraction blanks. The R^2^ values for standard curves ranged from 0.984 to 0.995, with an estimated efficiency ranging from 84.2 to 95.1%. Littoral and pelagic eDNA concentrations varied substantially by lake (Table 3). After weighing for lake zone area, mean eDNA concentrations ranged from 592 copies/L in Cobb to 7805 copies/L in Olive.

**Table 3:** Lake zone area and corresponding eDNA concentrations (minimum and maximum observed eDNA concentrations per lake zone included in parentheses).

| Site | Pelagic area (ha) | Littoral area (ha) | Mean Pelagic eDNA (copies/L) | Mean Littoral eDNA (Copies/L) | Weighted Mean eDNA (Copies/L) |
| --- | --- | --- | --- | --- | --- |
| Cobb | 1.0 | 1.3 | 253.8 (39.6 - 557.9) | 854.6 (35.9 - 2650.4) | 592.2 |
| Dog | 8.5 | 3.1 | 3447.1 (683.8 - 9148.1) | 9796.7 (3705.6 - 16839.3) | 5131.1 |
| Helen | 1.2 | 1.3 | 1342.4 (854.3 - 1586.9) | 3514.4 (2083.2 - 5060.5) | 2445.9 |
| Margaret | 14.4 | 3.6 | 791.9 (706.8 - 968.1) | 3034.1 (814.3 - 5689.7) | 1240.4 |
| McNair | 0.7 | 1.0 | 2395.4 (2214.9 - 2495.9) | 3505.0 (3181.1 - 4886.4) | 3050.5 |
| Mud | 4.7 | 2.6 | 399.3 (261.7 - 580.7) | 1550.6 (628.3 - 3833.3) | 797.5 |
| Olive | 0.5 | 1.2 | 8084.6 (5115.9 - 11758.9) | 7684.7 (1839.6 - 11829.1) | 7805.1 |
| Ross | 4.6 | 2.0 | 790.5 (439.7 - 1101.9) | 1209.8 (3763 - 2576.0) | 917.4 |
| Temple | 1.6 | 1.7 | 1180.1 (854.3 - 1685.7) | 1850.3 (1133.6 - 3887.0) | 1530.6 |
| Hidden | 11.8 | 2.6 | 342.0 (149.3 - 472.4) | 2652.9 (1277.2 - 5758.1) | 847.2 |

Linear models for each density metric demonstrated positive and significant correlations with eDNA particle concentration (Table 4, Figure 3). Individual density, biomass density, and ASM accounted for 59%, 63%, and 78% of the variation in observed eDNA particle concentration (adjusted R^2^), respectively. AIC values indicated that individual density and biomass density metrics provided roughly equivalent model fit; however, the ASM metric provided substantially improved model fit relative to individual density and biomass density (ΔAIC of 5.7 and 4.6, respectively). Trends did not substantially change when littoral area per lake was halved (Table S1). CPUE did not exhibit a significant correlation with eDNA particle concentration (Table 4, Figure S4).

**Table 4:** Model results evaluating the relationship between eDNA particle concentration and density (fish/ha), biomass (kg/ha), allometrically scaled mass (ASM/ha), and CPUE.

| Model | F-value | P-value | Adj. $R^2$ | Log Likelihood | AIC | ΔAIC |
| --- | --- | --- | --- | --- | --- | --- |
| Density | 12.37<sub>(1,7)</sub> | 0.010 | 0.59 | -77.78 | 161.6 | 5.7 |
| Biomass | 14.76<sub>(1,7)</sub> | 0.006 | 0.63 | -77.26 | 160.5 | 4.6 |
| ASM | 29.4<sub>(1,7)</sub> | 0.001 | 0.78 | -74.95 | 155.9 | - |
| CPUE | 1.92<sub>(1,7)</sub> | 0.208 | 0.10 | -81.27 | 168.5 | 12.6 |

**Figure 3:**
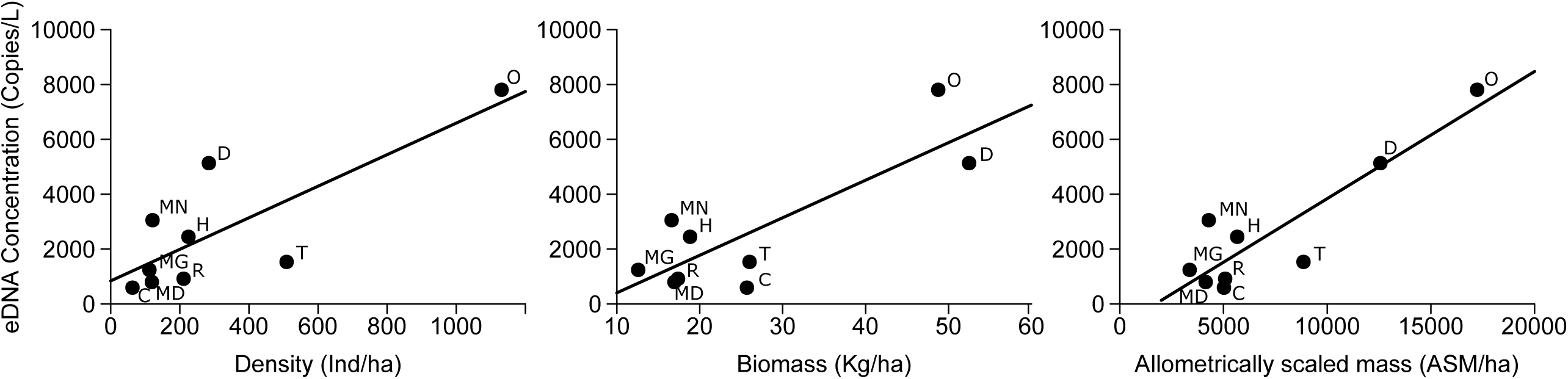
Correlation between weighted mean lake brook trout eDNA particle concentration and three metrics of abundance in the nine study lakes: (a) individual density (individuals/ha, R^2^ = 0.59), (b) biomass density (kg/ha, R^2^ = 0.63), and (c) allometrically scaled mass (ASM/ha, R^2^ = 0.78) (n = 9). O = Olive, D = Dog, C = Cobb, T = Temple, H = Helen, MD = Mud, MG = Margaret, MN = McNair, R = Ross.

### Estimating density and biomass from predicted allometrically scaled mass: a case study for population management

The eDNA concentration of Hidden Lake littoral and pelagic eDNA samples averaged 2653 and 342 copies/L, respectively, with a weighted mean average eDNA particle concentration of 847 copies/L (Table 3). Based on a linear model using data from the nine study lakes, Hidden Lake had an estimated ASM/ha of 4279.6 (Figure 4). After 1000 iterations, the mean number of individual mass values sampled from the Olive size distribution was 278.4, which represents the individual density (ind/ha) point-estimate for Hidden Lake; this corresponds to a total population size estimate of 3286 individuals. Predicted total biomass was 143.0 kg, with a biomass density of 12.1 kg/ha. Notably, point estimates of biomass density rank Hidden Lake lower than all nine study lakes, likely as a result of previous fish removal efforts between 2011 and 2017 in Hidden Lake. Upper 95% prediction intervals for population size, total biomass, density, and biomass density were 7629 individuals, 332.0 kg, 646.5 fish/ha, and 28.1 kg/ha, respectively. Due to the overall low concentration of eDNA present in the lake, lower 95% prediction intervals overlapped with zero for all four parameters.

**Figure 4:**
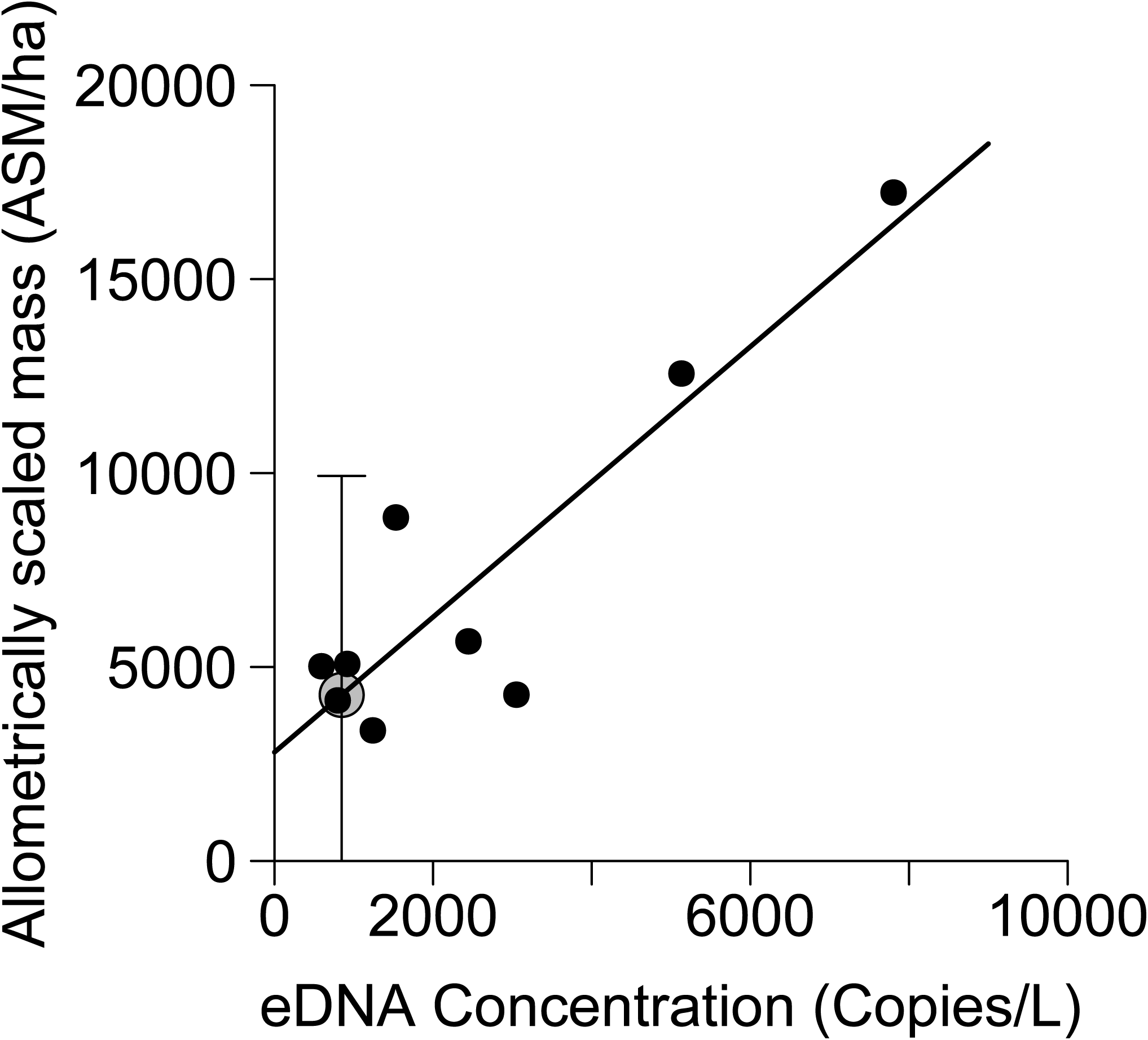
Predicting allometrically scaled mass (ASM/ha) for Hidden Lake based on eDNA particle concentration. Black dots represent values for the nine study lakes, gray circle represents the ASM/ha point estimate for Hidden Lake. Error bars represent 95% prediction intervals (n = 9).

### Predicting the allometric scaling coefficient for eDNA production in brook trout

Based on model AIC values, a scaling coefficient of 0.72 best explained patterns of eDNA particle concentration across the nine study lakes; models with scaling coefficients between 0.47 and 0.89 generated ΔAIC values < 2 (Figure 5). The ‘optimal’ scaling coefficient appeared to be slightly sensitive to the fraction of the area of each lake assigned to the littoral zone: when littoral zone area within each lake was halved, a scaling coefficient of 0.63 best explained patterns of eDNA particle concentration (Figure S5). However, credible intervals between the two models substantially overlapped; models with scaling coefficients between 0.28 and 0.84 generated ΔAIC values < 2 when lake littoral area was halved.

**Figure 5:**
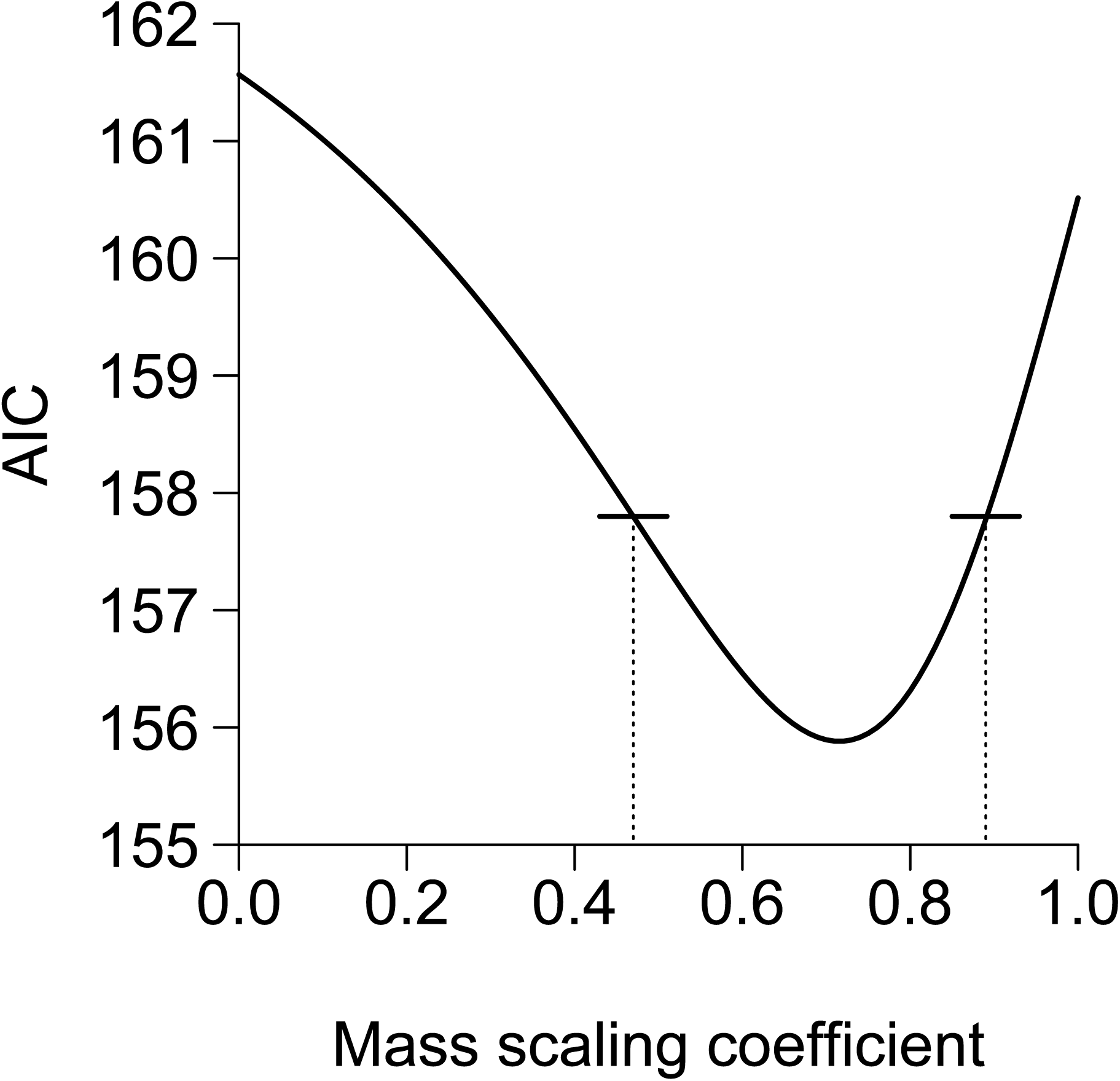
AIC values for models correlating brook trout eDNA and allometrically scaled mass (ASM), utilizing allometric scaling coefficients ranging from 0.00 (corresponding to individual density) to 1.0 (corresponding to biomass density). Horizontal black bars and dotted lines denote range of models with ΔAIC < 2 relative to the ‘optimal’ scaling coefficient (0.72).

## Discussion

Our study provides strong support for the hypothesis that eDNA production scales non-linearly with mass according to a power function. Incorporating allometric scaling coefficients to account for the distribution of biomass across individuals substantially improved predictive models, indicating that the distribution of biomass across individuals within a population may have an important effect when scaling individual eDNA production rates to the population-level. Incorporating metabolic scaling coefficients for mass into models of eDNA particle concentration and organismal abundance may therefore be particularly important in species that exhibit substantial inter-population variation in size distributions. Our findings contribute to a broader understanding of the ecology of eDNA production and have important implications for many eDNA applications. While the focus of this study was on the relationship between eDNA particle concentration and abundance using qPCR techniques, allometry in species with variable size structure could, for example, partially account for the variation observed in read numbers across environments in metabarcoding studies.

This study also reaffirms previous findings that metrics of population abundance correlate with species-specific eDNA particle concentration in natural environments (Klobucar, Rodgers, & Budy, 2017; Nevers et al., 2018; Pilliod et al., 2013; Schmelzle & Kinziger, 2016; Thomsen et al., 2012). Previous research has demonstrated a moderate correlation between density and/or biomass and eDNA particle concentration in lotic systems for brook trout (Baldigo et al., 2017; Wilcox et al., 2016). We found similar relationships within lentic systems, but also demonstrate that they can be considerably improved by integrating allometric scaling coefficients into estimates of organismal abundance. Notably, in eight of the nine study lakes the mean concentration of eDNA observed in lentic zone samples was higher compared to pelagic zone samples. eDNA particle concentrations generally show a strong correlation with the spatial distribution of fish within a lake (Ghosal et al., 2018; Hanfling et al., 2016), and our findings reflect well documented ecological preferences of brook trout, which tend to favor littoral zones (Magnan & Fitzgerald, 1982; Tiberti et al., 2017). The only lake where this trend was not observed was Olive, where pelagic and littoral zone eDNA concentrations were similar; this lake was also the smallest (and shallowest, at 3.5 m maximum depth) lake with the highest individual density of brook trout, indicating that fish are likely relatively evenly distributed across the lake.

The correlation coefficients we observed between eDNA concentration and all three metrics of abundance were greater than most previous studies conducted in nature (Yates et al., 2019). The relatively strong correlations we observed between our abundance metrics and eDNA concentration could also be due to the methodology with which we assessed population size. Our estimates of population size were obtained using mark-recapture studies and unbiased measures of size-structuring, which provided precise and standardized estimates of individual density, biomass density, and ASM. However, such estimates are rare in published eDNA/abundance studies; conducting mark-recapture studies to estimate population size is time consuming and requires a substantial commitment of labour and resources. To date only a handful of eDNA studies in nature have specifically enumerated population size (Klobucar et al., 2017; Levi et al., 2019; Tillotson et al., 2018) rather than proxies for abundance, such as CPUE (Yates et al., 2019). CPUE may be appropriate if it exhibits a strong correlation with abundance, but in some systems CPUE can perform poorly as a proxy for abundance (Hubert et al., 2012; Rose & Kulka, 1999). In our study systems CPUE did not exhibit a significant correlation with individual density and, as a result, eDNA concentration. Some of the substantial unexplained variation in nature between eDNA concentration and abundance observed in other systems could result from reliance on CPUE as a ‘proxy’ for abundance, although we acknowledge that for many species it may often be impractical or impossible to directly estimate population size.

It is important to note, however, that our abundance estimates may miss a small fraction of the adult population and do not account for juvenile (age 0+) abundance because fish were not included in the mark-recapture study until they were at least 80mm (to avoid excessive tagging mortality). Population size estimates therefore represent underestimates of true population census size. Discrepancies in juvenile abundance/density across lakes could account for some of the remaining unexplained variation present in our model, particularly since smaller fish would be expected to exhibit higher mass-specific eDNA production rates. Similarly, temperature is known to have a strong effect on metabolic rates (Brown et al., 2004) and eDNA production (Jo et al., 2019). Notably, bioenergetics models for a closely related species (bull trout, *Salvelinus Confluentus*) demonstrate that both the value of the normalization constant (*I*_0_) as well as the allometric scaling coefficient (*b*) can change with temperature (Mesa, Weiland, Christiansen, Sauter, & Beauchamp, 2013). Temple lake exhibited a substantially lower concentration of eDNA than expected from its ASM estimate; at 3.5 □, Temple lake was also substantially colder than the other eight study lakes during eDNA sampling (8.9-17.2 □). Although we lacked the replication to do so, integrating other important environmental variables (e.g. temperature, pH, etc.) into models of eDNA particle concentration across environments could further improve predictive models.

Despite these caveats, we demonstrate that it is possible to predict estimates of population abundance with eDNA samples and size structure data in similar ecosystems that lack abundance data. We predicted traditional metrics of abundance for Hidden Lake based on a hypothetical assumption that size structure in Hidden Lake closely resembled size structure in another study lake (Olive lake). Although predicted density metrics for Hidden Lake based on an ‘proxy’ Olive Lake size structure distribution were low and exhibited wide upper 95% prediction intervals, they still provided enough information to facilitate relative comparisons to the nine study lakes; we can predict with some certainty, for example, that if size structure in Hidden Lake was similar to that found in Olive, it would have had a lower biomass density relative to two of the nine study lakes (Dog and Olive). Furthermore, 95% prediction intervals represent a relatively stringent criteria of certainty; 75% or 80% prediction intervals might still represent useful information to help guide managerial or research decisions, although that would be up to practitioner discretion.

Most significantly, our results highlight the need for further empirical studies exploring and validating allometric scaling via power functions as a framework for modelling eDNA particle production rates. While we demonstrate that incorporating allometric scaling coefficients substantially improves models predicting abundance and eDNA concentration at the population level, we have not directly quantified how eDNA production scales allometrically in brook trout at the level of individual organisms. Nevertheless, recent experiments demonstrate that mass-specific eDNA production rates tend to decline as individual mass increases (Maruyama et al., 2014; Mizumoto et al., 2018; Takeuchi et al., 2019). We found that a scaling coefficient of 0.72 best described patterns of eDNA concentration for our study species across our nine study lakes; this value is closely aligned with the metabolic scaling coefficient for brook trout from (Hartman & Cox, 2008). Scaling coefficients between 0.51 and 0.87 produced models with ΔAIC values < 2; we therefore predict that the ‘true’ allometric scaling coefficient for eDNA production in brook trout will likely fall within this interval, although we do note that this point estimate was slightly sensitive to the area of each lake assigned to the littoral zone. If the area assigned to the littoral zone of each lake is halved, the value of the ‘optimal’ scaling coefficient is reduced to 0.63 (models with ΔAIC values < 2 range from 0.28-0.84) which is closer to theoretically expected values for excretory/consumptive/shedding allometric scaling coefficients (although note that credible intervals for both values substantially overlap). In future studies, detailed bathymetry data would be useful to disentangle these issues. Nevertheless, credible intervals for both models overlapped substantially, indicating that allometric scaling substantially improved explanatory models. To validate our findings, test our subsequent predictions, and disentangle what processes are likely to most strongly affect the value of eDNA scaling coefficients (e.g. metabolism vs excretion/shedding), further experiments are necessary to quantify allometric scaling of eDNA production at the individual level in brook trout.

As a well-supported general theory in ecology, experimental designs developed to test MTE hypotheses (e.g. (Allegier et al., 2015; Hartman & Cox, 2008)) can inform future experiments examining the effect of allometry on eDNA production rates. Notably, previous experiments investigating allometric scaling in excretion or metabolic rates quantified rates at the level of *individual* organisms (Allegier et al., 2015; Hartman & Cox, 2008; Vanni & McIntyre, 2016). Previous laboratory experiments quantifying the effect of biomass on eDNA production/shedding rates typically pooled organisms to create different biomass treatments (Doi, Uchii, Takahara, & Matsuhashi, 2015; Klymus et al., 2015; Lacoursière-Roussel, Rosabal, & Bernatchez, 2016; Mizumoto et al., 2018; Takahara et al., 2012). At best, such experiments pool organisms from similar size-classes, in which case eDNA production/abundance relationships across ‘treatments’ only reflect changes in abundance within a specific age- or size-class. Such experimental designs are likely to produce a strong relationship between eDNA concentration and biomass, as has been found in a meta-analytic review (Yates et al., 2019). While such studies were necessary to empirically quantify a preliminary correlation between eDNA particle concentration and metrics of abundance, they might obscure critical differences in mass-specific eDNA production rates across size classes that could have important consequences for population-level rates. Natural populations often exhibit substantial variation in the distribution of body size across individuals; the failure to account for allometric scaling in the relationship between biomass and eDNA production might partially explain the failure to translate the strong relationships observed in laboratory experiments to nature (Sebens, 1987). Notably, our eDNA/abundance models utilizing ASM exhibited correlation coefficients comparable to those typically observed in laboratory environments (Yates et al., 2019).

It may be possible to investigate allometry in eDNA production by pooling individuals that are the same size within replicates. However, we would advise against this because behavioural interactions between fish at high density in confined spaces may impact eDNA production; some studies have demonstrated that eDNA production per fish increases at high densities (Id et al., 2019). Brook trout, for example, are known to exhibit aggressive behaviour towards conspecifics (McNicol, Scherer, & Murkin, 1985), which could increase eDNA particle concentration at high densities due to increased activity and/or injuries inflicted upon each other. If size classes exhibit different behaviour at high densities, this could further affect estimates of allometric scaling. Future studies examining allometric scaling in eDNA production should therefore incorporate individuals from a gradient of age/size classes and quantify organismal eDNA production at the *individual*-level, as in (Takeuchi et al., 2019). Notably, the two studies to examine eDNA production rates at an individual level across age/size classes found that larger, older individuals exhibited lower mass-specific eDNA production rates (Maruyama et al., 2014; Takeuchi et al., 2019). There is also a critical need to conduct such experiments *in situ* at field study sites on wild organisms, as in (Pilliod, Goldberg, Arkle, & Waits, 2014). Laboratory experiments, while important from a validation perspective, may not reflect natural excretion processes because study organisms are housed in artificial conditions, fed artificial diets, and are often subject to fasting regimes (Vanni & McIntyre, 2016). Furthermore, size-scaling coefficients for metabolic processes such as nutrient excretion exhibit substantial interspecific variation and can even include values greater than 1 (Allegier et al., 2015; Vanni & McIntyre, 2016). Allometric scaling in eDNA production may therefore exhibit similar variability across species and should be investigated on a case-by-case basis.

Finally, our experiment investigated intraspecific allometry in eDNA production. Although there is substantial taxonomic variation, multiple studies have demonstrated that it is possible to extend allometric power-scaling across taxonomic groups for metabolic and/or excretory processes (Allegier et al., 2015; Vanni & McIntyre, 2016). Metabarcoding studies exhibit a weak but positive relationship between read count and organism biomass (Lamb et al., 2019). If allometric scaling in eDNA production exhibits a similar relationship across taxonomic groups, the relationship between read count and organism abundance could be strengthened by integrating allometry.

### Conclusions

Our results provide evidence supporting the hypothesis that eDNA production scales allometrically with organism mass We have demonstrated that the incorporation of additional (but straightforward to collect) size structure data to integrate key allometric scaling predications resulted in substantial improvement in models of eDNA concentration across environments. The bulk of experiments examining eDNA in nature have typically focused on presence/absence applications for species detection utilizing metabarcoding technologies (Goldberg et al., 2015), in which the detection of rare DNA fragments is often prioritized. As a result, substantial consideration in the literature has been given to factors that affect eDNA degradation and dispersion (e.g. (Barnes et al., 2014; Goldberg, Strickler, & Fremier, 2018; Harrison, Sunday, & Rogers, 2019; Strickler, Fremier, & Goldberg, 2015)), while relatively less attention has focused on the ecology of eDNA production. Our study demonstrates that the ecology of eDNA production may represent an understudied yet critically important subject, particularly when attempting to infer abundance from eDNA concentrations in nature. Future studies on eDNA/abundance relationships in nature should consider incorporating allometry, particularly when study species exhibit substantial inter-population variation in size distributions. However, there is also a need to validate this hypothesis in controlled experimental contexts at the level of individual organisms. As a well-developed ecological theory validated by numerous empirical studies (Vanni & McIntyre, 2016), the literature on the MTE represents a robust methodological foundation that future studies can utilize to explore relationships between a variety of environmental and ecological factors that might influence organismal production of eDNA. Such studies could further improve predictive models estimating abundance from eDNA particle concentration to the extent that, in some circumstances, species-specific eDNA particle concentration might be a reliable ecological indicator of abundance.

Predictive models would need to be calibrated on a system- and species-specific basis. The extent to which models for a particular species can be extended to different ecosystems or geographical regions also remains unknown. Future studies employing the methodology developed herein will likely need to construct models from population size/abundance estimates combined with standardized size distribution data on an individual species/system basis. These studies will also need to collect size distribution data, in addition to eDNA samples, to predict the density or biomass of organisms in similar ecosystems that lack abundance data. Direct estimates of allometric scaling coefficients for study species would also likely improve predictive models, although metabolic or excretory allometric scaling coefficients estimated in other empirical studies on the same (or closely related) species may represent useful starting points. In the absence of any other empirical data, the general scaling coefficient predicted by the MTE (0.75) may also suffice.

Depending on the species studied, obtaining robust population size estimates and individual size distribution data to calibrate initial models can often be difficult, labour intensive, and come with a substantial monetary cost. However, the benefits might be substantial – the idea that future researchers or managers might be able to obtain reasonable estimates of abundance from eight water samples and a small number of gill net sets is, from an ecologist’s perspective, exciting.

## Acknowledgements

We would like to thank Brent Brookes, Jacob Farkas, Tom Ridgeon, Natalie Dupont, Thaïs Bernos, Laura Bogaard, Haley Tunna, Ben Kelley, Mélia Lagacé, and Mathilde Tissier for their assistance conducting the mark-recapture experiments and collecting eDNA samples and Joanne Littlefair for her help in the laboratory. We would also like to thank Taylor Wilcox for his advice on working with the qPCR primer/probe set, as well as the associate editor and three anonymous reviewers whose comments greatly improved the manuscript. This research was funded by a Fonds de recherche du Québec - Nature et technologies (FRQNT) team grant (AMD, MEC, DJF) and a Natural Sciences and Engineering Research Council of Canada (NSERC) strategic project grant (DJF, JP, AMD). DMG was funded by the NSERC Strategic Project Grant awarded to DJF and co-applicants. MCY was funded by an NSERC EcoLac post□doctoral scholarship and FRQNT post-doctoral scholarship.

## Data Accessibility Statement

eDNA particle concentration data for each lake will be deposited in the Dryad Digital Repository upon acceptance.

## Author contributions

MCY collected eDNA samples and analyzed eDNA data. DG collected and analyzed mark-recapture and size structure data. Statistical analyses were conducted by MCY. MCY wrote the first draft of the manuscript, and all authors contributed substantially to subsequent drafts.

## Figure Captions

Figure S1: Map of the nine study lakes located in Alberta and British Columbia, Canada. Figure S2: Timing of sampling activities in 2018. S.A. refers to size-structure assessment.

Figure S3: Relationship between catch-per-unit-effort (CPUE) of a large and small gill net and individual density (fish/ha) for the nine study lakes (adjusted R^2^ < 0) (n = 9).

Figure S4: Relationship between brook trout eDNA particle concentration and catch-per-unit-effort (CPUE) of a large and small gill net for the nine study lakes (R^2^ = 0.10).

Figure S5: AIC values for models correlating brook trout eDNA with littoral lake area halved and allometrically scaled mass (ASM), utilizing allometric scaling coefficients ranging from 0.00 (corresponding to individual density) to 1.0 (corresponding to biomass density). Horizontal black bars and dotted lines denote range of models with ΔAIC < 2 relative to the ‘optimal’ scaling coefficient (0.72).

## Notes

### Competing Interest Statement

The authors have declared no competing interest.

### Summary of Updates

Minor changes to analytical methodology, new analysis testing sensitivity of littoral area estimates, and further discussions of broad implications of findings. Also minor revisions to include sampling details and improve clarity.

