## Supplementary figures and images for "The relationship between eDNA particle concentration and organism abundance in nature is strengthened by allometric scaling"

### Figure S1

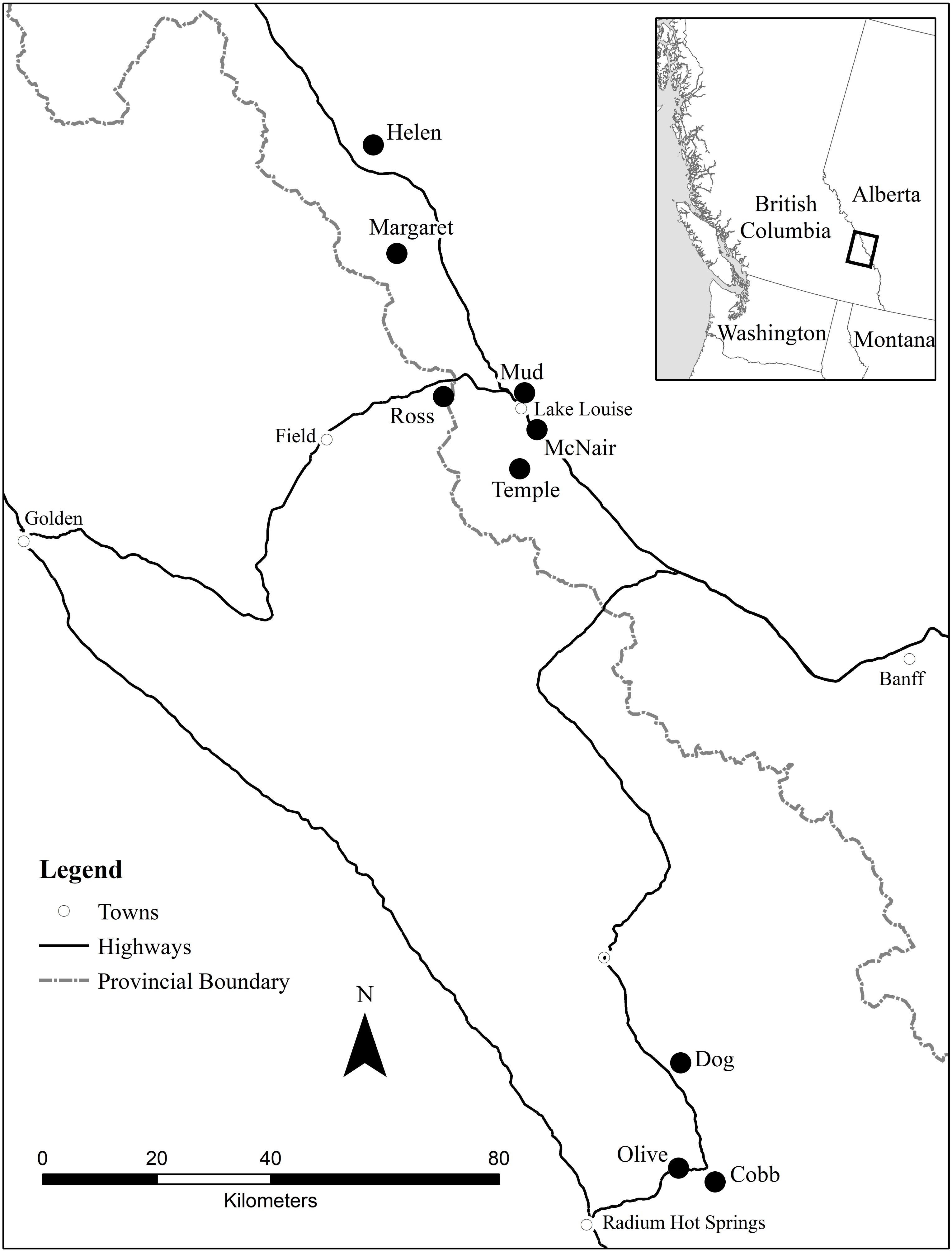

### Figure S2

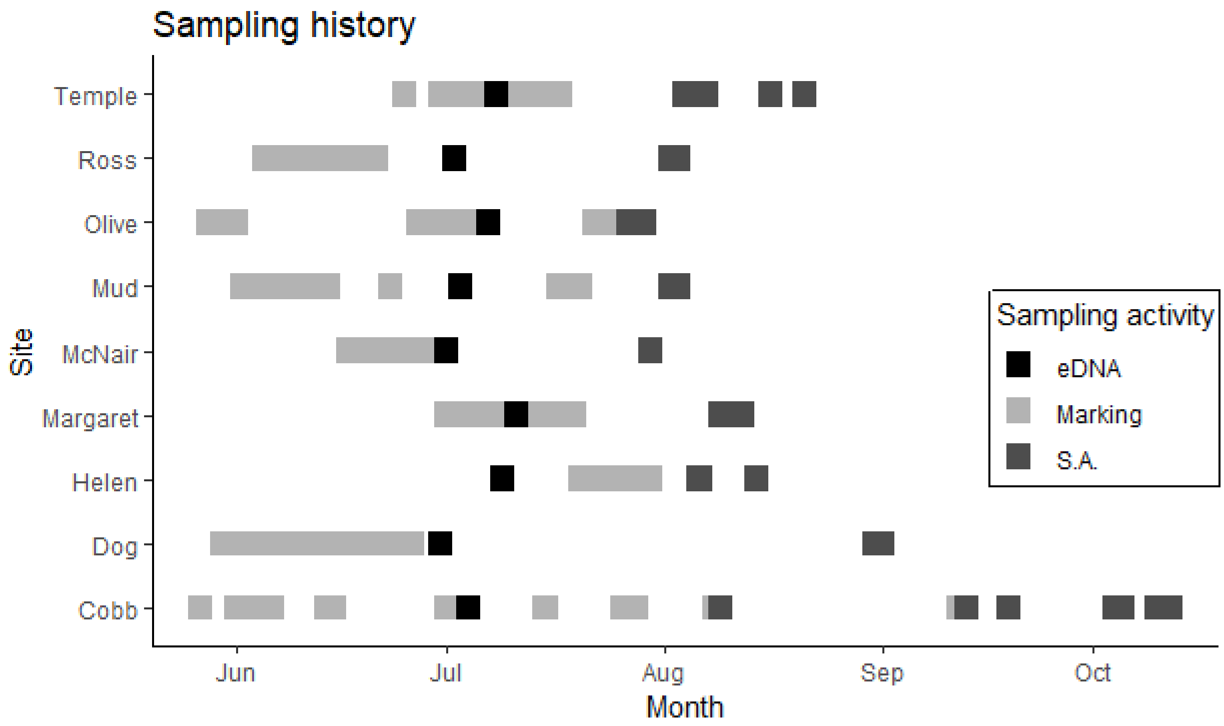

### Figure S3

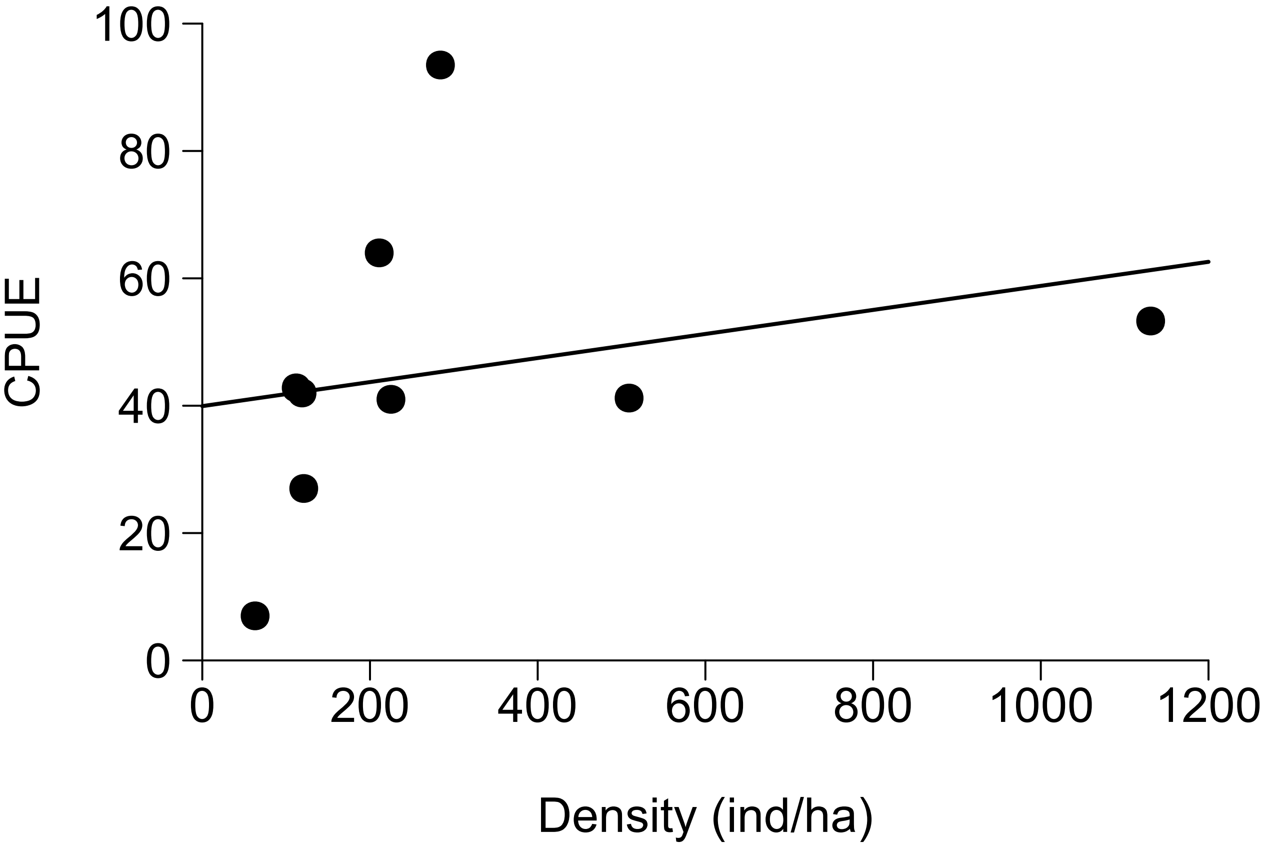

### Figure S4

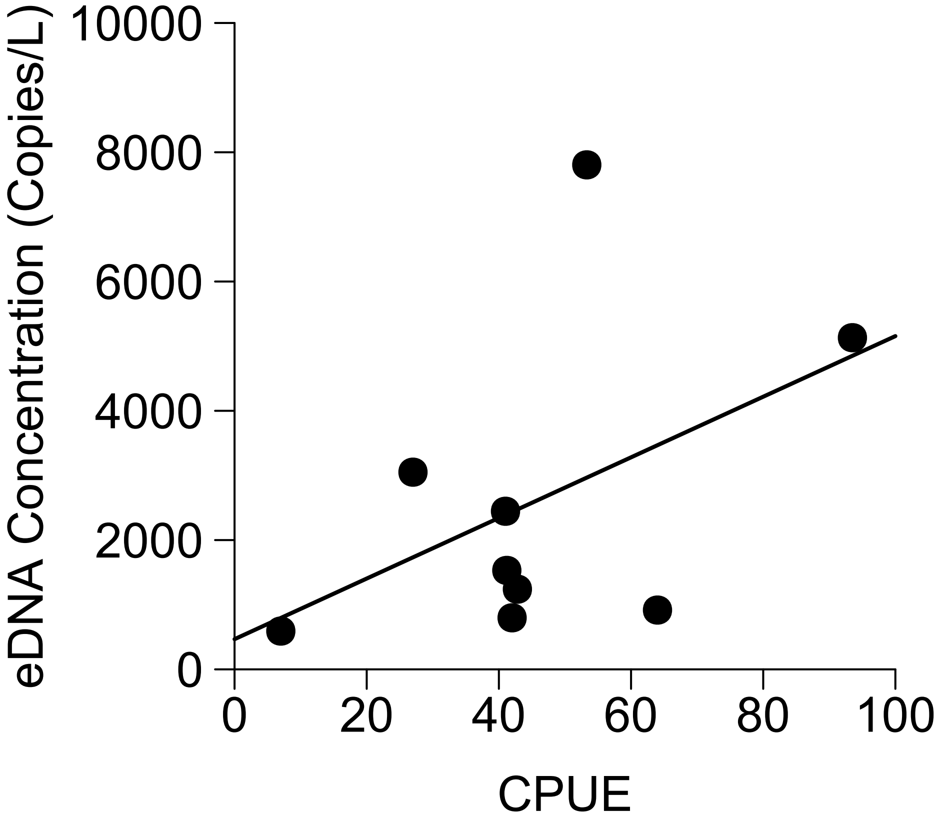

### Figure S5

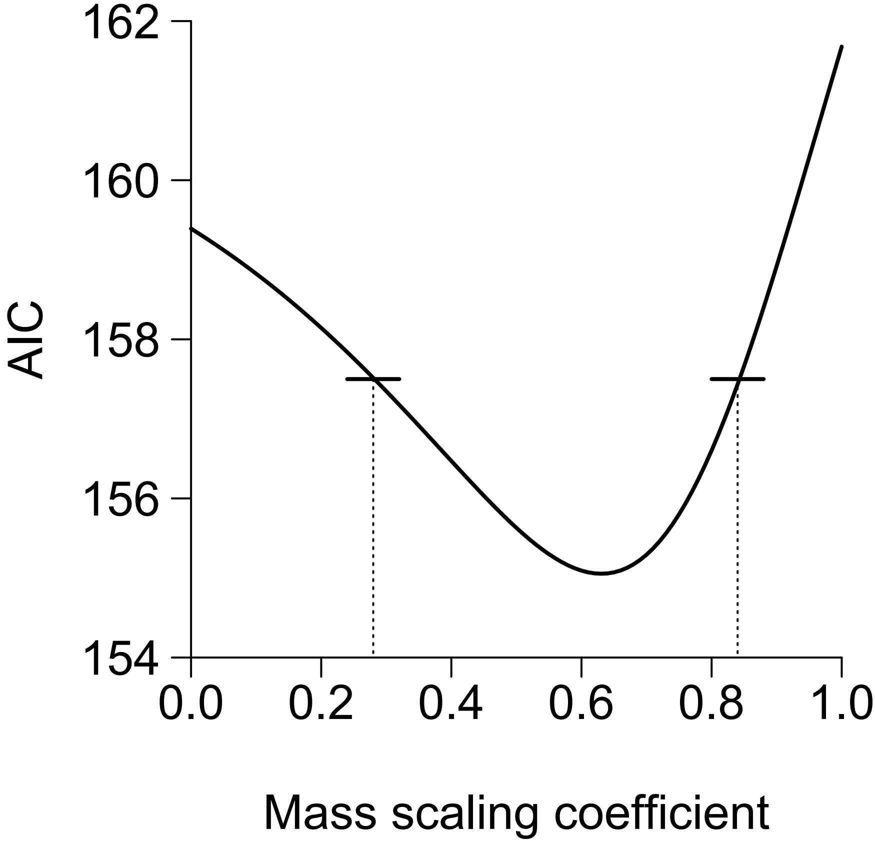
