## Appendix S1 for "The relationship between eDNA particle concentration and organism abundance in nature is strengthened by allometric scaling"

### Extracting eDNA (Qiagen DNeasy Mini kit) for eDNA applications from glass fibre filters

**Legend:**
Red = Important notes
Blue = optional

##### Important points before starting

- All centrifugation steps are carried out at room temperature (15–25°C) in a microcentrifuge.
- Proteinase K must be stored at 4°C upon receiving the kit.
- ALWAYS RUN AN EXTRACTION BLANK, for every set of extractions you do. This tests for contamination that may occur during extractions.

##### Things to do before starting

- Buffer ATL and Buffer AL may form precipitates upon storage. Warm both solutions to 56°C until the precipitates have fully dissolved. You might want to premake the AL:EtOH mixture (1:1)
- Buffer AW1 and Buffer AW2 are supplied as concentrates. Before using for the first time, add the appropriate amount of ethanol (96–100%) as indicated on the bottle to obtain a working solution.
- Heat the AL:OH buffer mixture before adding it to the samples, otherwise precipitate will form.
- Heat buffers AW1 and AW2, as well, and make sure you shake them vigorously. AW1 is the most important. Heating ensures that salts in the buffers are dissolved completely – samples where buffers are not heated may end up with substantial salt contamination in end products; this may inhibit downstream PCR applications.
- Preheat an incubator, thermomixer, shaking water bath, or rocking platform to 56°C for step 3.
- If using frozen samples, equilibrate the sample to room temperature. Avoid repeated thawing and freezing of samples since this will lead to DNA degradation.
- ALWAYS use filter tips when pipetting.
- Change gloves any time sample liquid gets on them – major potential source of contamination.
- DO NOT pipette fully to the "second stop" – this will create bubbles with a lot of the Qiagen kit reagents that bubble up over the lip of tube; this is a major source of potential contamination.

##### Procedure

1. Prepare ATL:Proteinase K mixture (made fresh daily). Pulse Vortex 5s.
    - Note: Filters already stored in ATL buffer in Yates et al. 2020.
2. Samples already have 700ul of ATL in each tube. Add 80ul of proteinase K to each tube. Try to distribute Proteinase K throughout the tube (on multiple filter sides, in liquid at top, in center of rolled filter, etc.). Vortex all tubes after adding proK for 10 seconds.
3. Ensure the caps are closed and labelled. Place the tubes in an incubator. Incubate at 56°C (it will become gelatinous). Incubate overnight. Vortex incubating tubes once approximately 3 hours after digestion begins.
4. After incubation, transfer filters (using decontaminated tweezers, see note below) to a Qiashredder tube and run lysed buffer and filter membrane through labelled Qiashredder column. MAKE SURE

YOU LABEL THE SIDE OF THE COLLECTION TUBE AS WELL. You will require ***TWO*** columns per sample; the whole filter + buffer WILL NOT fit into a single Qiashredder column. Elute lysis buffer for 2 minutes at 11,000 rpm.
Note: make sure that you have a 30% bleach/DI bath, an ultrapure water rinse, and two sets of tweezers prepared. You must decontaminate tweezers for at least 10 minutes between handling samples. Alternate tweezers used to move filters to Qiashredder columns so that one is in decontamination bath at all times to improve efficiency. Make sure to rinse decontaminated tweezers in ultrapure water bath – do not introduce bleach droplets directly into your samples.

5. After centrifugation, remove spin column and **CAP** Qiashredder collector tube (caps come with package). IMMEDIATELY vortex both columns. This is done to resuspend pellet at the bottom of the collector tube.

6. After vortexing, transfer elutant to new 5ml sterile microcentrifuge tube. DO NOT TRANSFER UNLESS YOU HAVE RESUSPENDED PELLET FROM STEP 5.

7. During incubation, prepare AL:EtOH mixture at 1:1 ratio (1400 µl of the mixture per sample). Ensure it is at around 56°C.
Note: Prepare AL:EtOH mixture first thing next morning after overnight incubation. For filters stored with 700ul, you will need to add 1400ul of AL:EtOH and mix in the large 5ml microcentrifuge tubes.

8. Add 1400µl of Buffer AL:EtOH mix to the sample (in 5ml centrifuge tubes), and mix thoroughly by vortexing. A white precipitate may form on addition of Buffer AL and ethanol. Warming the AL:EtOH mixture will reduce formation of precipitate.

9. Pipette the mixture from step 8 (including any precipitate) into the DNeasy Mini spin column placed in a 2 ml collection tube (provided). (Be careful not to cross-contaminate). Centrifuge at 6000 x g (8000 rpm) for 1 min. Empty collection tube, but DO NOT DISCARD COLLECTION TUBE – return spin column to empty collector tube. You will need to re-use the collector because the maximum capacity of a spin column is 600ul. DO NOT PUT MORE INTO A SPIN TUBE. You will need to empty collector tube and perform this step four+ times. Do not discard collector tube at the end of this step – you will need it in step 10.

10. Place the DNeasy Mini spin column in the emptied 2 ml collection tube from step 9, add 500 µl Buffer AW1, and centrifuge for 1 min at 6000 x g (8000 rpm). DO NOT discard collection tube, just empty the elutant; you will need to reuse the tube in step 11. Make sure you HEAT buffer AW1 before adding to spin column.

11. Place the DNeasy Mini spin column in the emptied 2 ml collection tube from step 10, add 500 µl Buffer AW2, and centrifuge for 3 min at 20,000 x g (14,000 rpm) to dry the DNeasy membrane. Discard flow-through and collection tube. Make sure you HEAT buffer AW2 before adding to spin column.
Note: Make sure to re-use spin column from step 9-10, you will need a clean spin column for the next step and to deposit eluted DNA in.

12. Place the DNeasy Mini spin column in a new collection tube. Centrifuge for 1 min at 20,000 x g (14,000 rpm) to dry the DNeasy membrane. Discard flow-through and collector tube. The function of this step is to remove any residual ethanol remaining from previous steps. Note that many of the collector tubes will have small droplets of liquid at the bottom after centrifugation – if you did not perform this step, that liquid would remain in your final eluted DNA product. For this application, we are eluting with VERY small quantities of AE buffer at the next (and final) step – that last droplet would represent a substantial proportion of the final total elutant volume if you did not remove it during this step. Note that ethanol is a KNOWN PCR INHIBITOR – you do not want this source of potential inhibition in your final elutant.
Note: Following the centrifugation step, remove the DNeasy Mini spin column carefully so that the column does not come into contact with the flow-through, since this will result in carryover of ethanol. If carryover of ethanol occurs, empty the collection tube, then reuse it in another centrifugation

13. Place the DNeasy Mini spin column in a clean 2 ml collector tube and pipet 65 µl Buffer AE directly onto the DNeasy membrane. Incubate at 56°C for 5 min, and then centrifuge for 1 min at 6000 x g (8000 rpm) to elute.
    ** You can elute with TE or AE.
    Alternatively: If application is extremely sensitive to any form of contamination (e.g. metabarcoding), use sterile DNA/RNA/DNase/RNase free 1.5 microcentrifuge tube with the cap removed in place of collector tubes.

14. On the same DNeasy Mini spin column and collector tube as step 13 pipet 65 µl Buffer AE directly onto the DNeasy membrane. Incubate at 56°C for 5 min, and then centrifuge for 1 min at 6000 x g (8000 rpm) to elute.

15. Using pipette and filter tips, transfer extracted DNA solution from collector tube to pre-labelled sterile DNA/RNA/DNase/RNase free 2ml microcentrifuge tube for long-term storage.
    Note: MAKE SURE you pipette the final eluted liquid up and down several times in collector tube prior to transferring to 2ml microcentrifuge tube. This is done to resuspend any pellet that may have formed from the centrifugation during the elution stages (steps 13 and 14)
